# Establishment of a pig influenza challenge model for evaluation of monoclonal antibody delivery platforms

**DOI:** 10.1101/2020.03.12.988808

**Authors:** Adam McNee, Trevor Smith, Barbara Holzer, Becky Clark, Emily Bessell, Ghiabe Guibinga, Heather Brown, Katherine Schultheis, Paul Fisher, Stephanie Ramos, Alejandro Nunez, Matthieu Bernard, Veronica Martini, Tiphany Chrun, Yongli Xiao, John C. Kash, Jeffery K. Taubenberger, Sarah Elliott, Ami Patel, Peter Beverley, Pramila Rijal, David Weiner, Alain Townsend, Kate Broderick, Elma Tchilian

## Abstract

Monoclonal antibodies are a possible adjunct to vaccination and drugs in treatment of influenza virus infection. However questions remain whether small animal models accurately predict efficacy in humans. We have established the pig, a large natural host animal for influenza, with many physiological similarities to humans, as a robust model for testing monoclonal antibodies. We show that a strongly neutralizing monoclonal antibody (2-12C) against the hemagglutinin head administered prophylactically at 15 mg/kg reduced viral load and lung pathology after pandemic H1N1 influenza challenge. A lower dose of 1 mg/kg of 2-12C or a DNA plasmid encoded version of 2-12C, reduced pathology and viral load in the lungs, but not viral shedding in nasal swabs. We propose that the pig influenza model will be useful for testing candidate monoclonal antibodies and emerging delivery platforms prior to human trials.

---

Influenza virus infection remains a significant global health threat to humans and livestock causing substantial mortality and morbidity. Monoclonal antibodies (mAbs) administered either prophylactically or therapeutically have been proposed as a strategy to provide immediate immunity and augment existing vaccines and drugs in combatting seasonal and pandemic influenza infection. Broadly neutralizing antibodies against conserved epitopes of the haemagglutinin (HA) stem and head, and antibodies against the neuraminidase (NA) are candidates for human treatment ^1, 2^. Both prophylactic and therapeutic administration of these antibodies have been shown to be effective in the mouse and ferret^3–10^. However, early results from human clinical trials showed that efficacy in mice and ferrets is not always predictive of outcome in humans^11–14^. The reasons for the apparent lack of efficacy in humans are not clear but may include the difficulty of achieving high serum and nasal concentration in a large body mass, the potency of the mAbs or the challenge of therapeutic administration in the face of a high viral load. Variability due to pre-existing immunity in human experimental or natural infection challenge studies is an additional problem. There is therefore a need for a large animal model in which mAbs selected on the basis of *in vitro* assays and efficacy in small animals can be further studied to help in selecting promising mAbs and determining how best to administer them in clinical trials. Pigs may provide such a model. They are large animals and a natural host for influenza viruses. Pigs and humans are infected by the same subtypes of virus, have the same distribution of sialic acid receptors in their respiratory tract and are physiologically, anatomically and immunologically more similar to humans than small animals^15, 16^.

While great progress in antibody delivery is being made, the high costs that are associated with the production, purification and quality control are major challenges in the development of clinical mAbs against influenza and other infectious diseases. Moreover, long-term protection is difficult with a single inoculation because of the short half-life of the mAb. Alternative *in vivo* antibody gene transfer strategies using DNA, RNA or viral vectors have shown that antibody genes can be stably maintained in the host tissue, resulting in potent and long-term expression of mAbs in the body following a single administration^17–23^. DNA plasmid encoded mAbs (dMAbs), which are delivered to muscle tissue, are a novel approach with the potential to provide durable immunity^24–26^. The plasmid DNA is well-tolerated and nonintegrating, does not require cold-chain distribution, can be delivered repeatedly and is relatively inexpensive to produce. Previous studies have demonstrated the efficacy of such an approach for protection against influenza in mice ^27^.

We have previously tested therapeutic administration of the broadly neutralizing anti-stem FI6 antibody in the pig influenza model. This did not reduce viral load in nasal swabs and bronchoalveolar lavage (BAL), although there was reduction of pathology after aerosol delivery^28^. Broadly neutralizing anti-stem mAbs are less potent at direct viral neutralization as compared to anti-head antibodies and require Fc receptor engagement for *in vivo* protection^29, 30^. We demonstrated that human IgG1 FI6 did not bind to pig Fc receptors, perhaps accounting for the weak effect. Therefore to establish a more robust pig model we reasoned that a strongly neutralizing strain specific anti-head HA mAb should overcome this problem and give clear protection, providing a benchmark against which other mAbs and delivery platforms might be tested. Here we used the 2-12C mAb isolated from an H1N1pdm09 exposed individual, which shows strong neutralizing activity and selects influenza virus variants with HA substitutions K130E^31^. Furthermore we administered 2-12C prophylactically to provide the best chance to reveal an effect on viral load, as it is challenging for therapeutic administration to reduce viral load after infection has been established. We went on to evaluate the potential of an *in vivo* produced 2-12C using synthetic dMAb technology in support of the translation of this technology to humans as an immunoprophylactic.

## Results

### Establishment of a prophylactic pig influenza challenge model with recombinant 2-12C mAb

To establish a positive control protective mAb and delivery method in the pig influenza model we tested the strongly neutralizing human IgG1 anti-head 2-12C mAb^31^ in a prophylactic experiment. We administered 15 mg/kg of 2-12C intravenously and the control groups received an isotype matched IgG1 mAb or the diluent. Twenty-four hours later the pigs were challenged with pandemic swine H1N1 isolate, A/swine/England/1353/2009 (pH1N1) and 4 days later culled to assess viral load and pathology (**Fig. 1a**). 2-12C significantly decreased viral load in nasals swabs on each day over the course of infection, although the decrease was less on days 2 and 3 than days 1 and 4 **(Fig. 1b)**. The total viral load in nasal swabs over the 4 days in animals treated with 2-12C was also significantly less as determined by the area under the curve (AUC) compared to diluent and isotype controls (p=0.0267 and p=0.021). Viral load in the 2-12C group was significantly reduced in the BAL and no virus was detected in the lungs at 4 days post infection (DPI). Overall 2-12C had a clear effect on viral load in nasal swabs, BAL and lung.

**Figure 1.**
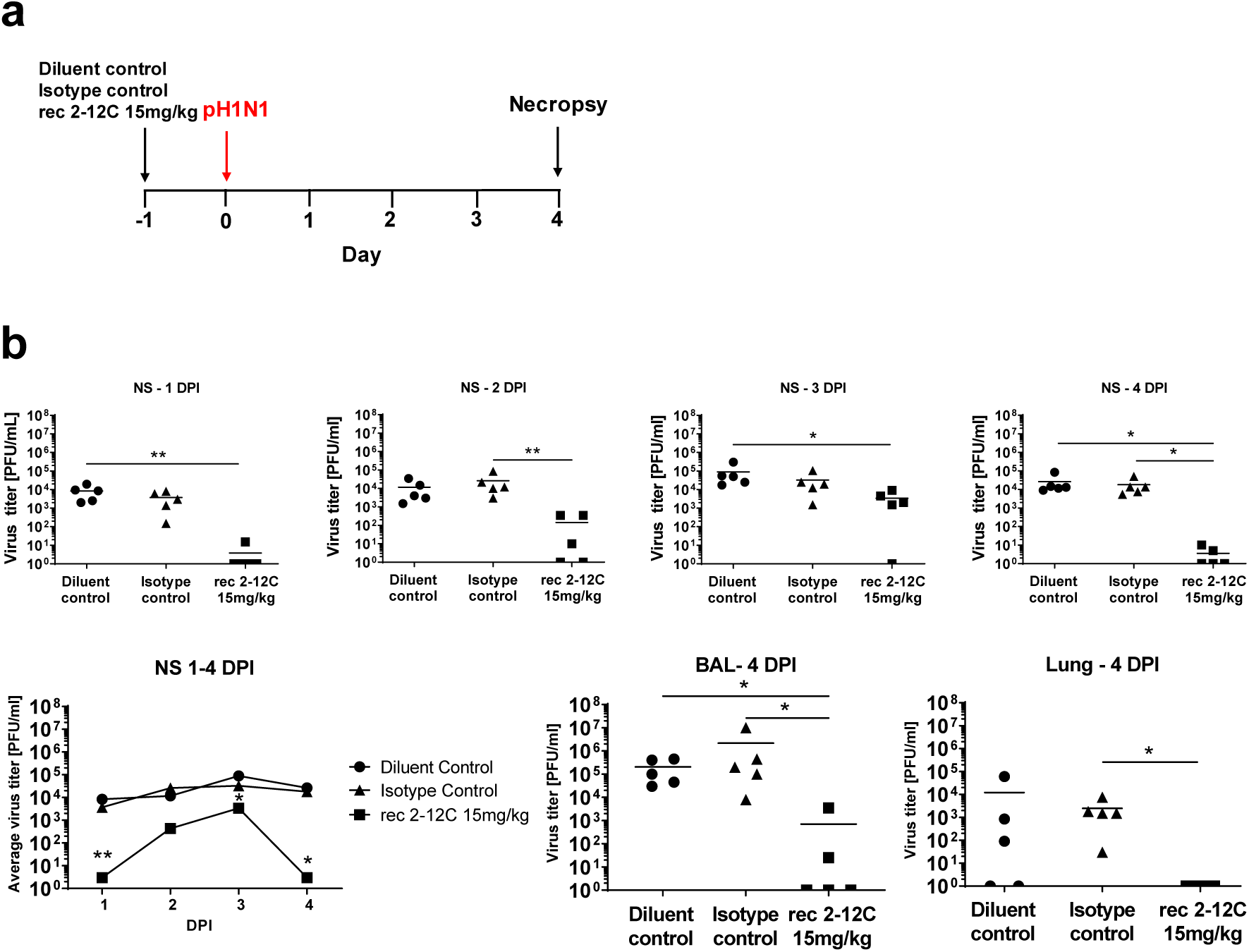
Experimental design and viral load. Antibodies or diluent were administered intravenously to pigs, which were infected with pH1N1 virus 24 hours later. Nasal swabs (NS) were taken at 1, 2, 3 and 4 days post infection (DPI) and pigs sacrificed at 4 DPI **(a)**. Viral titers in NS, accessory lung lobe (Lung) and BAL at 4DPI were determined by plaque assay **(b).** Each data point represents an individual pig within the indicated group and bars show the mean. Viral shedding in NS was also represented as the mean of the 5 pigs on each day and significance versus diluent control indicated by asterisks. Viral titers were analysed using one-way non-parametric ANOVA, the Kruskal-Wallis test. Asterisks denote significant differences *p<0.05, **p<0.01, versus indicated control groups.

The isotype and diluent control groups showed the most severe gross pathology and histopathology scores (**Fig. 2a**). The red tan areas of pulmonary consolidation were present mainly in the cranial and middle lung lobes of animals from the isotype and diluent groups, but no lesions were found in the 2-12C group **(Fig. 2b)**. Similarly, necrotizing bronchiolitis and bronchial and alveolar exudation, and mild thickening of the alveolar septa was shown in the animals from both control groups. Minimal to mild alveolar septa thickening and lymphoplasmacytic peribronchial infiltration was present in only one animal from the 2-12C treated group. Influenza A nucleoprotein (NP) (brown labelling) was detected by immunohistochemistry (IHC) in the bronchial and bronchiolar epithelial cells, alveolar cells and exudate in the bronchioles and alveoli in both control groups. No NP labelling was found in the lung of animals treated with 2-12C.

**Figure 2.**
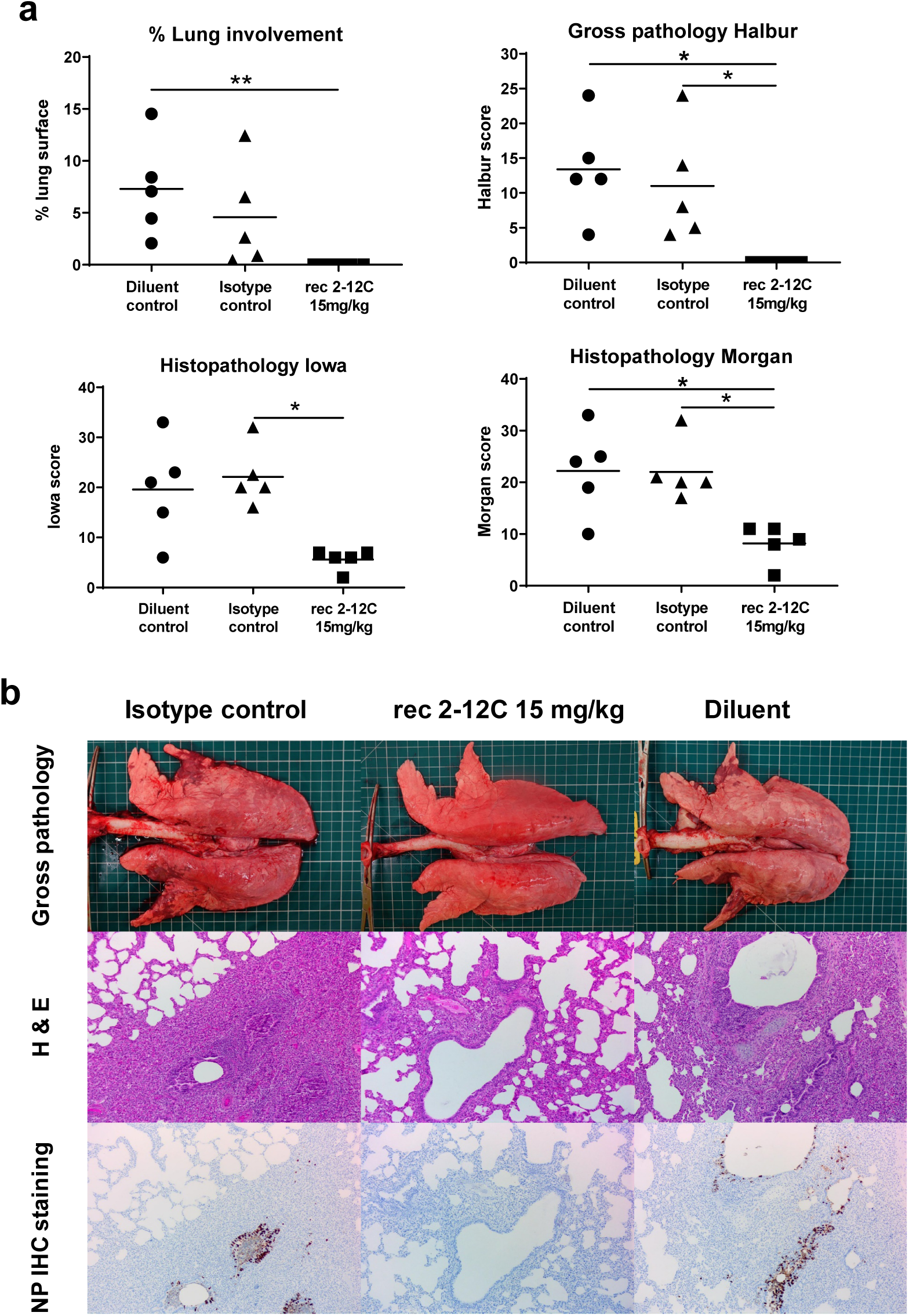
Lung pathology. Antibodies or diluent were administered intravenously to pigs, which were infected with pH1N1 virus 24 hours later. The animals were culled at 4 DPI and lungs scored for appearance of gross and histopathological lesions. The score for each individual in a group and the group means are shown **(a).** Representative gross pathology, histopathology (H&E staining; 100x) and immunohistochemical NP staining (200x) for each group are shown **(b).** Pathology scores were analysed using one-way non-parametric ANOVA with the Kruskal-Wallis test. Asterisks denote significant differences *p<0.05, **p<0.01, versus indicated control groups.

Recombinant 2-12C was detected in the serum at a mean concentration of 74.6 μg/ml and in the BAL at 183.9 ng/ml at 4 DPI as determined by an HA ELISA **(Fig. 3a)**. HA specific IgG was also detected in nasal swabs at a mean concentration of 24 ng/ml. The serum of the 2-12C treated group exhibited strong neutralizing activity at a 50% inhibition titer of 1:8,000 and 1:164 in the BAL **(Fig. 3b)**. Neutralizing activity in nasal swabs could not be tested due to limited sample availability. These data show that prophylactic intravenous administration of recombinant 2-12C at 15 mg/kg significantly reduced viral load and pathology in the pig, a large natural host animal.

**Figure 3.**
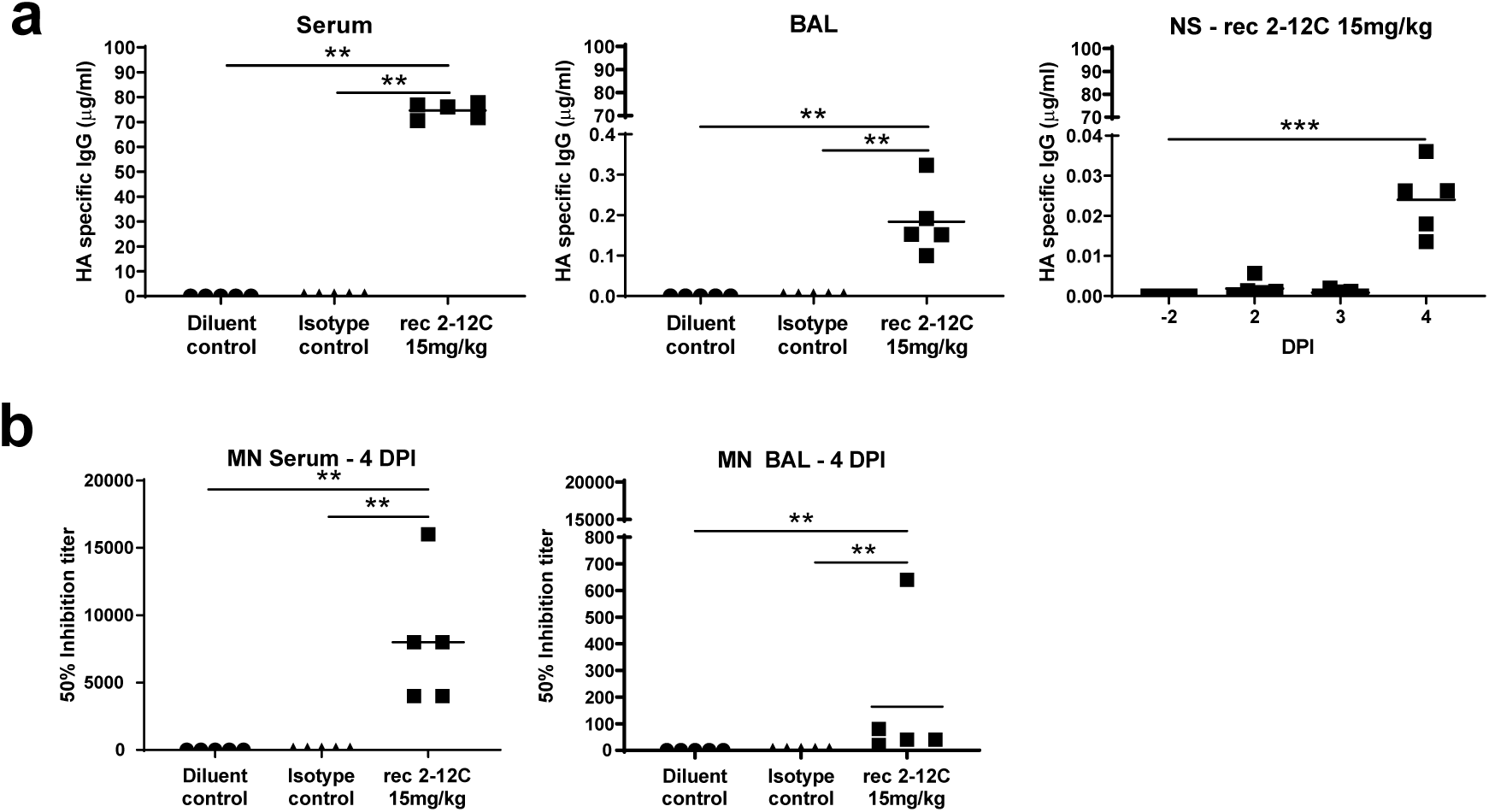
Concentration and neutralising titre of 2-12C in serum and mucosal tissues. H1 HA specific IgG in serum, BAL at 4 DPI and nasal swabs (NS) at the indicated DPI **(a).** 50% neutralization titers against pH1N1 in the serum and BAL at 4 DPI **(b).** Symbols represent an individual pig within the indicated group and lines the mean. Data were analysed using one-way non-parametric ANOVA with the Kruskal-Wallis test. Asterisks denote significant differences *p<0.05, **p<0.01 and ***p<0.001, versus indicated control groups.

### Design and expression of DNA encoded mAb (dMAb) in pigs and mice

To investigate the potential of a DNA-launched influenza mAb to mediate disease protection in the pigs we designed and engineered dMAb 2-12C (**Fig. 4a**). The gene sequences of the human IgG1 heavy and light chains were codon and RNA-optimized and inserted into a single modified-pVax1 DNA expression vector plasmid, separated by furin and P2A peptide cleavage sites. Expression of dMAb 2-12C was confirmed by Western Blot analysis, the band was detected at the same molecular weight as rec 2-12C (**Fig. 4b**). To assess *in vivo* expression, dMAb 2-12C was formulated with human recombinant hyaluronidase, as an optimization to enhance gene expression in the context of delivery with adaptive *in vivo* electroporation^32^. We measured expression of human IgG in myocytes 3 days after the administration of dMAb 2-12C pDNA into the tibialis anterior muscle of BALB/c mice, confirming local expression of human IgG at the site of delivery (**Fig. 4c**). To assess levels and durability of *in vivo* expression, mice were conditioned to reduce the T cell compartment using CD4 and CD8-depleting antibodies, this has been shown to permit the expression of a human mAb construct in mice in the absence of a host anti-drug antibody (ADA) response^25^. Administration of dMAb 2-12C pDNA to the tibialis anterior muscle was associated an increased durability of circulating levels of the mAb in the serum compared to rec 2-12C, administered intravenously (**Fig. 4d**).

**Figure 4.**
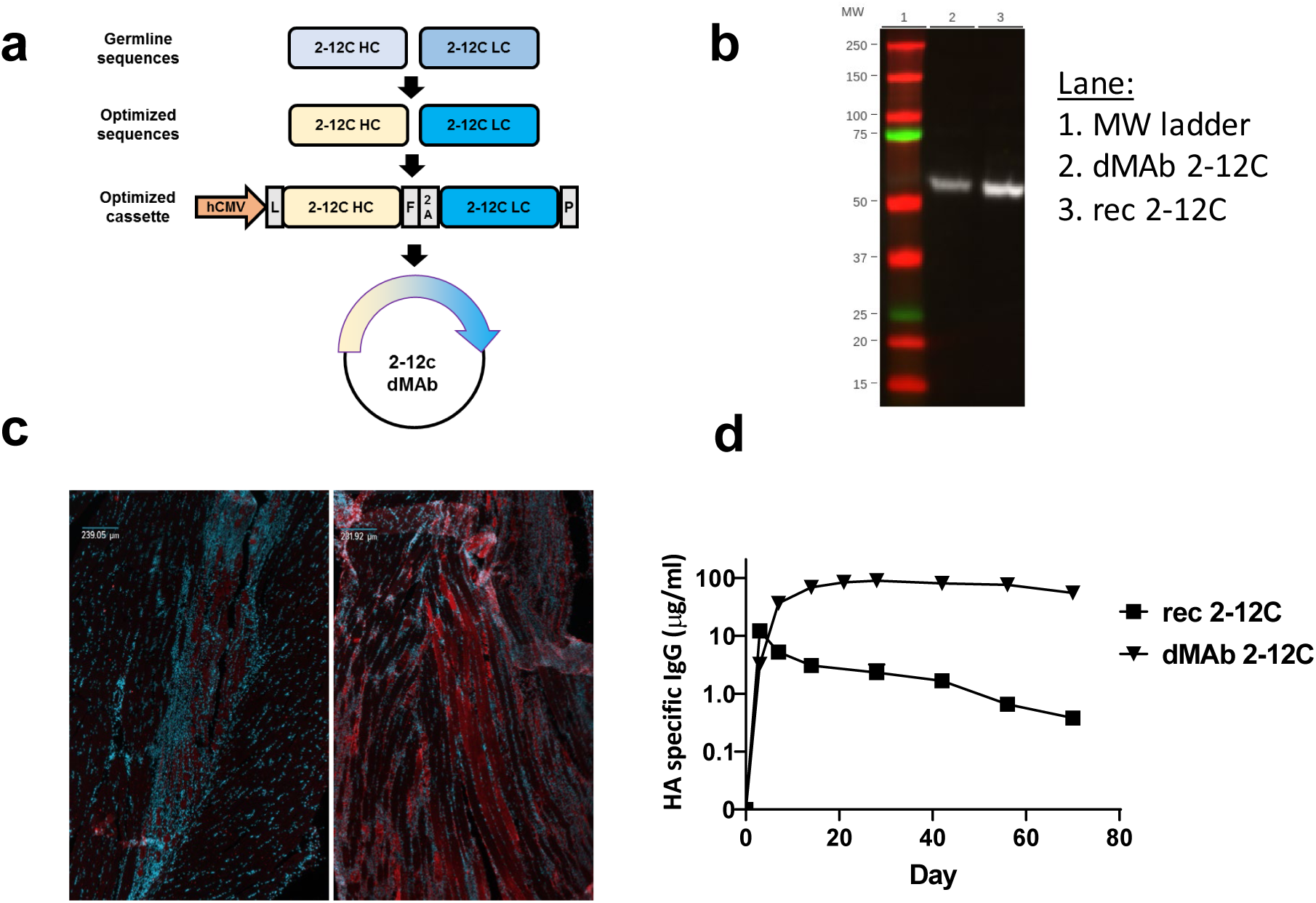
Design and expression of anti-influenza dMAb 2-12c. dMAb 2-12C single construct was designed to express human Ig light and heavy chain sequences codon and RNA-optimized for *in vivo* expression **(a)**. The optimized cassette includes a human CMV promoter (hCMV), a human IgG signal sequence (L), 2-12C HC and LC separated by a furin (F) and P2A cleavage site, and bovine growth hormone polyadenylation signal (P). *In vitro* expression of dMab 2-12C **(b)**. Supernatant from dMAb 2-12C plasmid transfected 293 T cells was run on a western blot next to recombinant 2-12C mAb. *In vivo* expression of dMAb 2-12C **(c, d)**. dMAb 2-12C or pVAX pDNA was administered intramuscularly by electroporation to BALB/c mice. Immunofluorescence images were taken on muscle tissue sections harvested 3 days after pDNA administration **(c)**. Blue = DAPI, red = human IgG. Serum human IgG levels after day 0 administration of recombinant or dMAb 2-12C in BALB/c mice quantified by ELISA **(d)**.

*In vivo* expression of dMAb 2-12C was measured in pigs. Expression of human IgG in myocytes 3 days after the administration of dMAb 2-12C pDNA into the quadriceps muscle of the pig, confirmed the local expression of the human IgG at the site of delivery (**Fig. 5a**). We measured serum levels of the expressed mAb after the day 0 administration of a total dose of 24 mg dMAb 2-12C pDNA into the quadriceps of the pig. Circulating levels of the expressed mAb in the serum were detected by an HA ELISA (**Fig. 5b**). Peak levels were 7 and 12 µg/ml for the two animals. Influenza H1N1 neutralizing activity (1:640 peak activity in both pigs) of the serum harboring the *in vivo* expressed 2-12C mAb was measured in a microneutralization assay (**Fig. 5c**). In both pigs, reductions in the serum levels of the expressed mAb and serum neutralizing activity were associated with the detection of an anti-drug antibody (ADA) response against the human 2-12C mAb (**Fig. 5b-d**). These results demonstrate that dMAb 2-12C induces expression of human 2-12C after intramuscular delivery by electroporation in mice and pigs.

**Figure 5.**
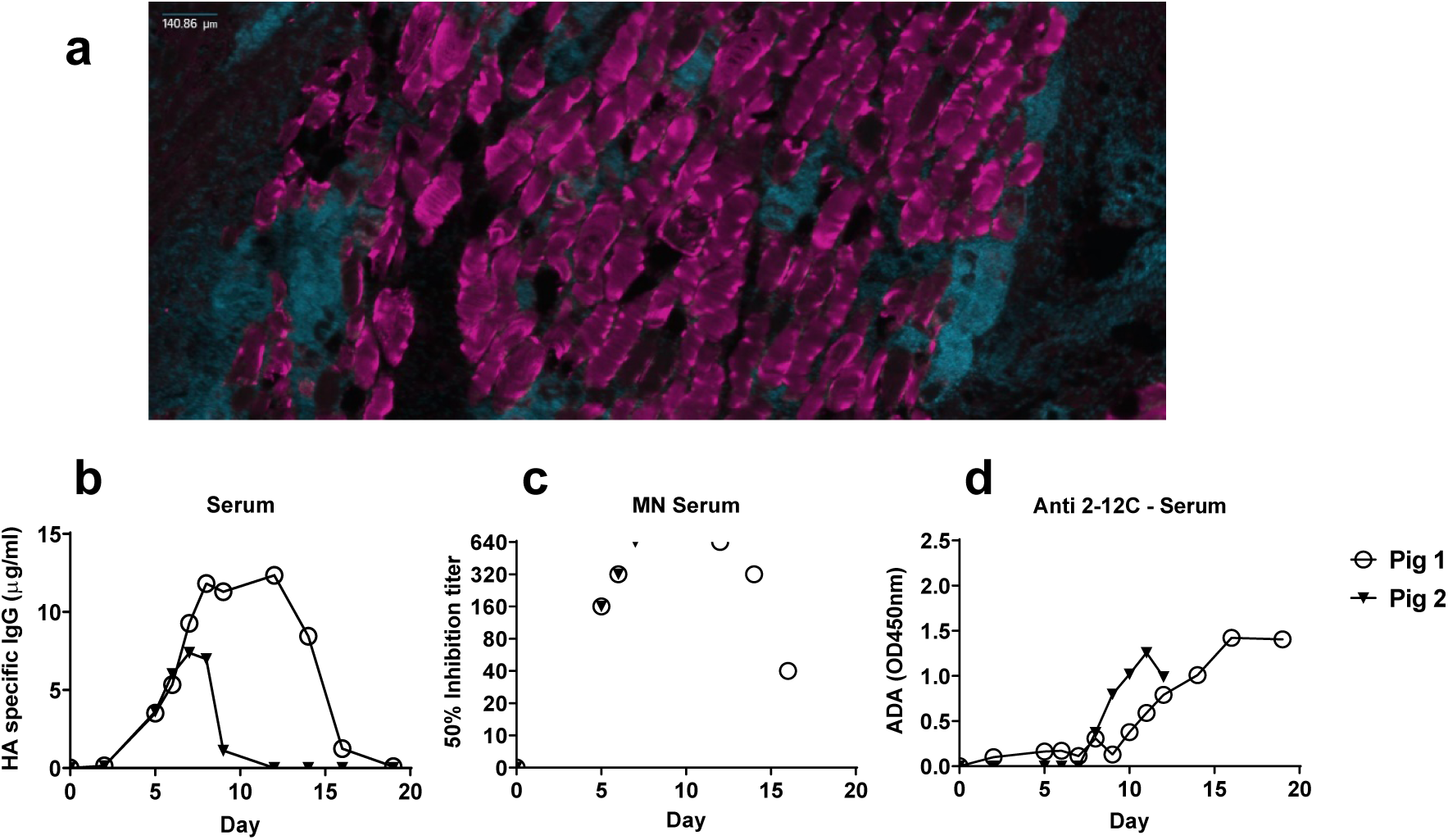
Expression of dMAb 2-12C in pigs. dMAb 2-12C pDNA was administered into the quadriceps muscle of Yorkshire pigs on day 0. Immunofluorescence images were taken on muscle tissue sections harvested 3 days after pDNA administration **(a)**. Blue = DAPI, red = human IgG. Serum 2-12C levels after day 0 administration of dMAb 2-12C were quantified by ELISA **(b)**. Serum neutralizing activity of H1N1 in a microneutralization assay **(c)**. Anti-drug-antibody (ADA) response against 2-12C mAb measured by ELISA **(d)**.

### Assessment of rec 2-12C dosing and dMAb 2-12C in a pig influenza model

After we had established 2-12C as a robust positive control antibody, we tested a lower dose of rec 2-12C and the efficacy of dMAb 2-12C in the pig influenza challenge model. The dMAb 2-12C was administered 6 days before the pH1N1 challenge to allow for accumulation of the *in vivo* expressed 2-12C mAb in the host. Preliminary experiments indicated the serum levels of 2-12C were generally not negatively impacted by the ADA until day 9, so we believed this schedule allowed a window of 3-4 days to test the efficacy of dMAb 2-12C. Recombinant 2-12C at 15 mg/kg and 1 mg/kg were delivered intravenously 24 hours before challenge to 6 pigs per group (**Fig. 6a**).

**Figure 6:**
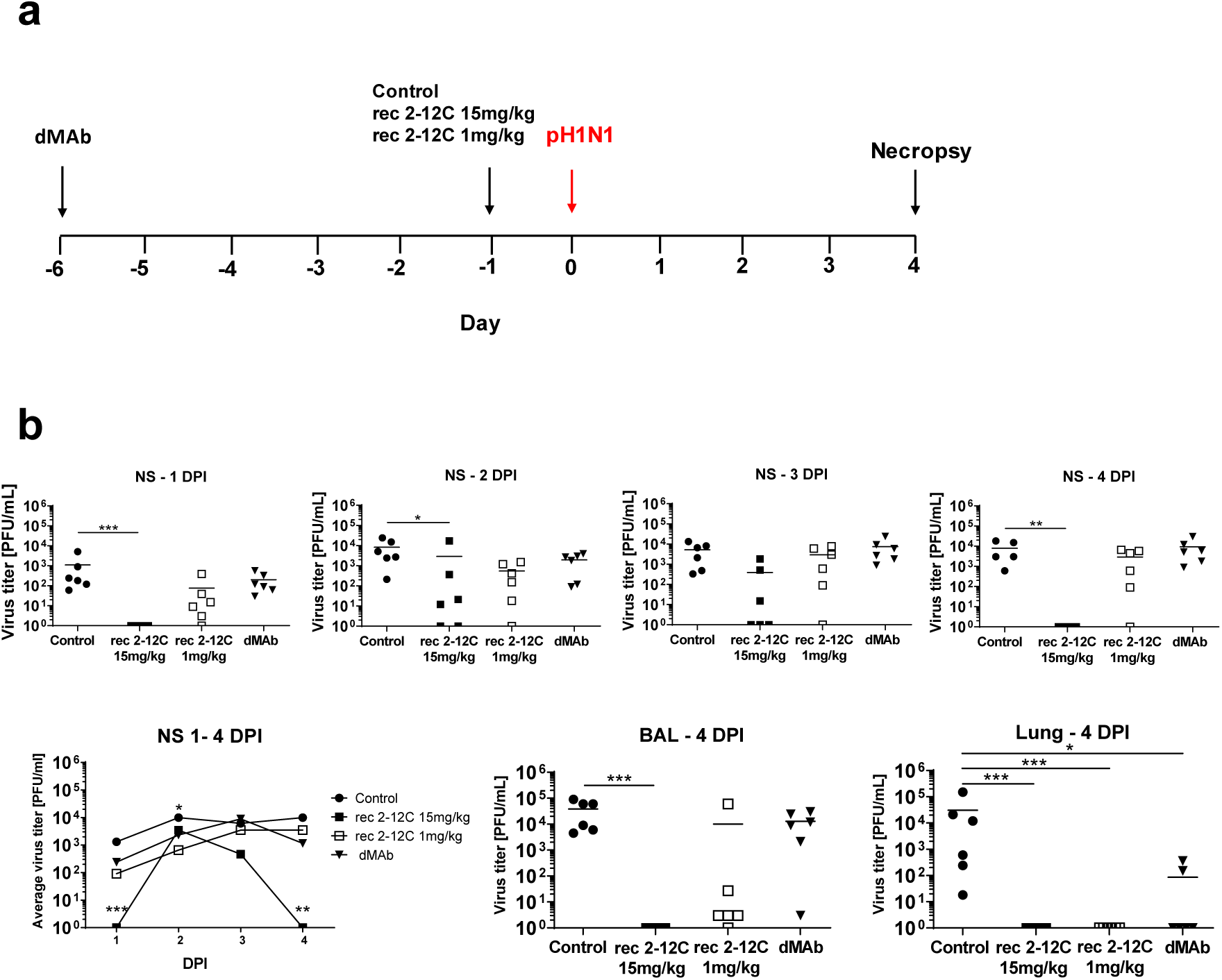
Experimental design and tissue viral load after recombinant and dMAb 2-12C treatment. Six days before pH1N1 challenge 6 mg of dMAb 2-12C was administered intramuscularly by electroporation. Recombinant 2-12C at 15 mg/kg and 1 mg/kg were delivered intravenously 24 hours before challenge. Control group was untreated animals. Nasal swabs (NS) were taken at 0, 1, 2, 3 and 4 DPI and pigs culled at 4 DPI **(a).** Viral titers in daily NS, BAL and accessory lung lobe (Lung) at 4 DPI were determined by plaque assay **(b).** Each data point represents an individual within the indicated group and bar is the mean. Viral shedding in NS is also shown as the mean of the 6 pigs over time and significance indicated against the control. Asterisks denote significant differences *p<0.05, **p<0.01 and ***p<0.001, versus control as analyzed by Kruskal-Wallis test.

As before, the greatest effect on virus replication in nasal swabs of 2-12C at 15 mg/kg was at day 1 and day 4 post infection, although no virus detected in BAL and accessory lung lobe **(Fig. 6b**). The AUC in the 2-12C group was significantly different from the control (p=0.0235). The lower dose of 2-12C at 1 mg/kg did not have a statistically significant effect on viral load in nasal swabs or BAL, although it reduced the titer in the lung accessory lobe. Similarly, dMAb 2-12C did not have a significant effect on viral load in nasal swabs or BAL, but significantly reduced viral load in lung.

The gross pathology was reduced in all experimental groups although this was significant only in the rec mAb groups, due to an outlier score in one animal in the dMAb 2-12C group. Histopathological evaluation also showed significantly decreased scores in all experimental groups (**Fig. 7a, b)**. All animals from the control group displayed changes consistent with a mild to moderate bronchointerstitial pneumonia, with various degrees of lymphohistiocytic septal infiltration and bronchial and perivascular cuffing, and with frequent areas of necrotizing and suppurative bronchiolitis. Viral antigen was detected by IHC in bronchial and bronchiolar epithelial cells, alveolar epithelial cells and exudate (macrophages) in all 6 control animals. Milder changes, mainly located in the septa, were found in animals from the antibody treated groups. The extent of bronchiolar changes and presence of IHC labelling varied in each group, although at least one animal per group displayed bronchiolar changes and virus antigen detection, in contrast to experiment 1. In addition to the differences in the histopathological scores, rec 2-12C 15 mg/kg group had 2 animals with acute bronchial lesions and antigen detection, whereas rec 2-12C 1 mg/kg group had 3 animals with bronchial lesions and antigen. The dMAb group had 1 animal with acute bronchial lesion and 5 animals where antigen was detected.

**Figure 7:**
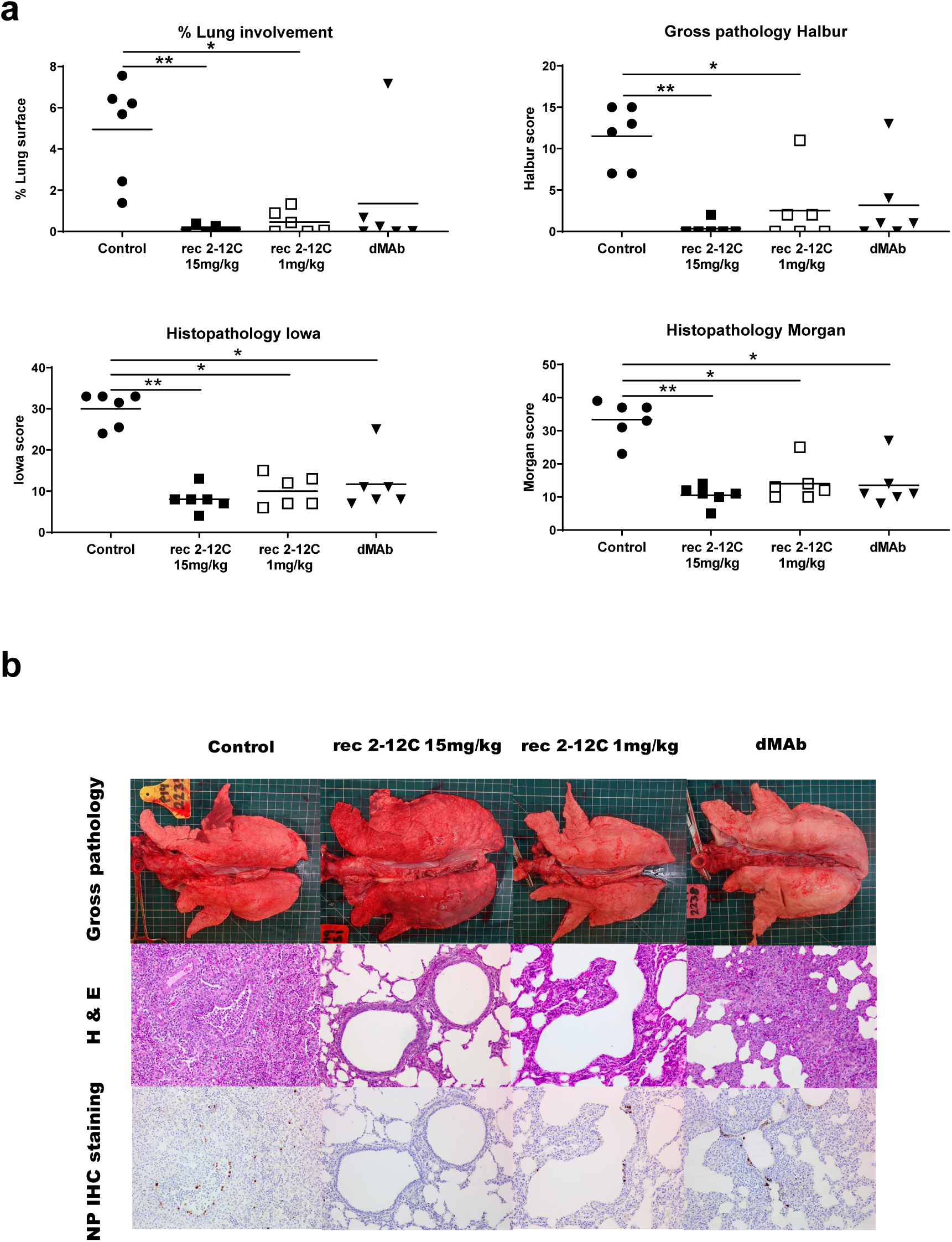
Lung pathology after recombinant and dMAb 2-12C administration. Six days before pH1N1 challenge 6 mg dMAb 2-12C was administered intramuscularly by electroporation. Recombinant 2-12C at 15 mg/kg and 1 mg/kg were delivered intravenously 24 hours before challenge. Controls were untreated animals. The animals were culled at 4 DPI and lungs scored for appearance of gross and histopathological lesions (**a).** Representative gross pathology, histopathology (H&E staining; 200x) and immunohistochemical NP staining (200x) for each group are shown **(b)**. Asterisks denote significant differences *p<0.05, **p<0.01 and ***p<0.001, versus control group as analyzed by Kruskal-Wallis test.

The concentration of the mAbs in the serum was determined daily after challenge. 2-12C was detected in the 15 mg/kg group at 101 μg/ml and in the 1 mg/kg group at 10 μg/ml 24 hours after administration. This declined in both groups over the next 3 days (**Fig. 8a**). In contrast the dMAb reached its peak of 0.99 μg/ml at day 7 after administration, declining thereafter. In the BAL HA specific antibodies were detected in the rec 2-12C groups (mean of 320.3 ng/ml for the 2-12C 15 mg/kg and 8 ng/ml for the 1mg/kg 2-12C groups) and a trace in the dMAb (1.5 ng/ml). In nasal swabs HA specific antibodies were detected in the 15mg/kg 2-12C group at 21.9 ng/ml 4DPI **(Fig. 8a)**.

**Figure 8:**
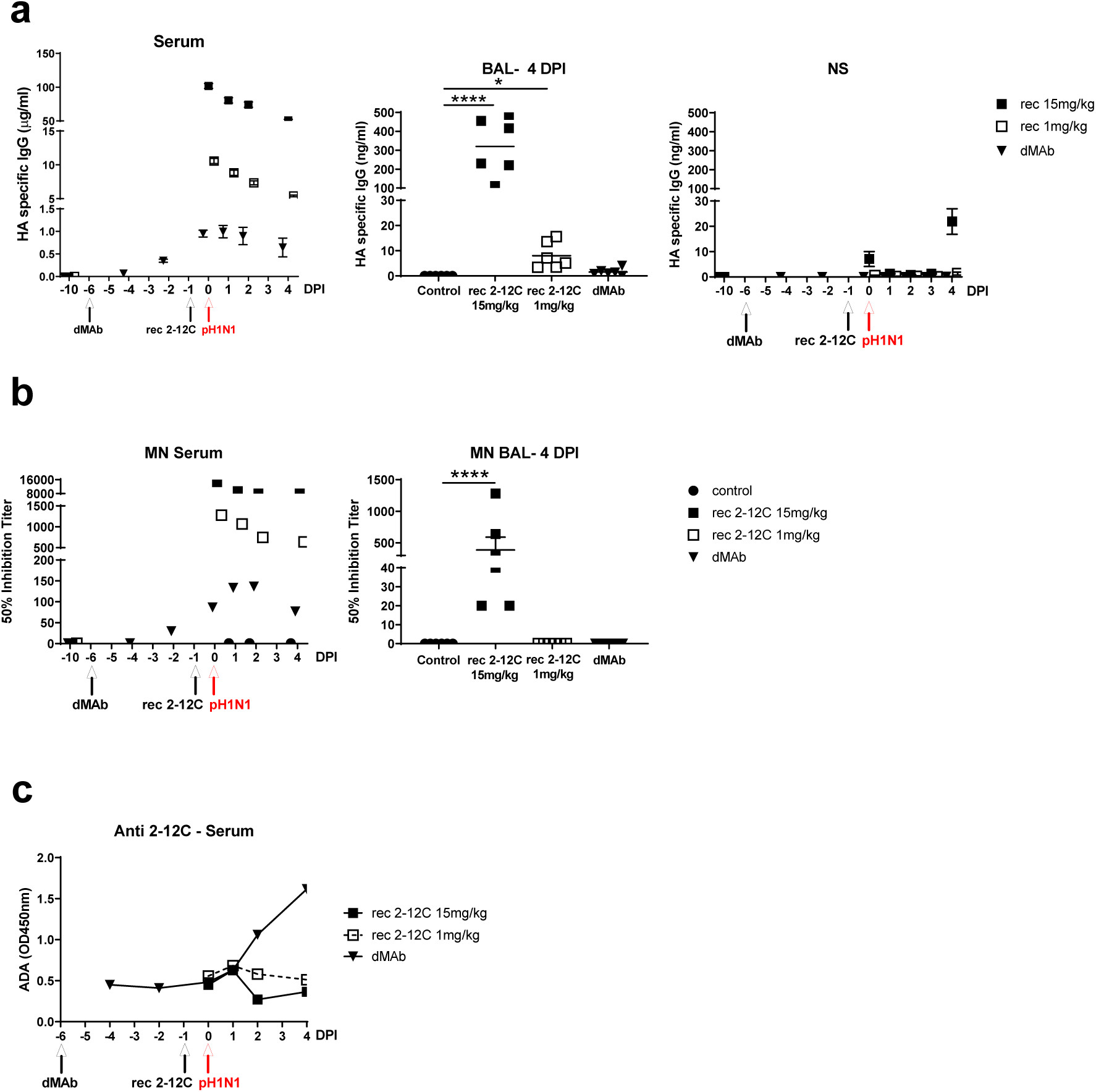
2-12C levels and anti-viral activity in serum and mucosal tissues after recombinant and dMAb 2-12C administration. The concentration of 2-12C in the serum, BAL and NS was quantified by HA specific ELISA at the indicated time points after administration and pH1N1 challenge **(a).** The mean 50% neutralization inhibition values for the individual groups in serum over time and BAL at 4 DPI **(b).** Anti-2-12C antibody (ADA) responses in serum detected by binding to immobilised 2-12C mAb **(c).** Data were analysed using one-way non-parametric ANOVA with the Kruskal-Wallis test. Asterisks denote significant differences *p<0.05, **p<0.01 and ***p<0.001, versus control group.

Neutralizing activity in serum was detected in all groups **(Fig. 8b)**. There was 50% inhibition titer of 1: 14,600 for the 15 mg/kg group and 1:1,280 for the 1mg/kg group at the time of challenge. A peak titer of 1:136 was detected at day 7 in the dMAb 2-12C-treated group. MN activity in BAL was only seen in the rec 2-12C 15 mg/kg group. A pig anti-2-12C ADA response was detectable in the dMAb group at day 8 after administration increasing at day 10 **(Fig. 8c)**. No ADA response was seen to the rec proteins in the serum, most likely because the animals were culled only 4 days after rec antibody administration.

Overall these results indicate that rec 2-12C offers robust protection at 15 mg/kg and that this correlates with mAb concentration and neutralization in serum. Administration of recombinant 2-12C protein at 1 mg/kg and dMAb 2-12C at 6 mg (0.5 mg DNA/kg) significantly reduced lung pathology and viral load in the lungs, but not in nasal swabs or BAL.

### Sequencing of virus

Inoculant virus and viruses from nasal swab samples of two control pigs and two 2-12C (15 mg/kg) treated pigs at 3 DPI were subjected to deep sequence analysis. In the inoculant, two control samples, and one of the 2-12C treated samples (53_2-12C), all segments of the influenza genome had 100% coverage with lowest average per base coverage of 98384.7 for NA segment in sample of 53-2-12C. For the other 2-12C treated sample (51_2-12C), the lowest segment coverage was 94.8% for the NA segment with average per base coverage of 2587.7. Although the inoculant had been passed in MDCK cells for at least 5 times, from these sequencing data, a total of only 5 SNPs (3 non-synonymous SNPs) were detected as compared to the reference A/swine/England/1353/2009 (pH1N1) sequence (Table 1). Non-synonymous SNPs found in sequenced nasal swab samples of 2 control and two 2-12C (15 mg/kg) treated pigs at 3 DPI compared to A/swine/England/1353/2009 are shown in Table 2. One non-synonymous SNP (HA K226E) that was detected in 2 control samples and one 2-12C treated sample (51_2-12C), was also identified as one of the HA SNPs found in the inoculant virus (Table 1). Three non-synonymous SNPs were detected in the other 2-12C treated sample (53_2-12C). One of them was also HA K226E, while the other 2 SNPs (HA G172E and HA T201I) were not detected in inoculant virus. Their frequencies in sample 53_2-12C are 12.1% (HA G172E) and 11.1% (HA T201I) respectively (Table 2). Therefore, from our deep sequencing data, we did not see evidence of escape of virus from antibody 2-12C (15 mg/kg) at day 3. No mutations in the 2-12C binding site were detected. Similarly no evidence for viral evolution driven by 2-12C was detected in day 4 nasal swab samples in the 1mg /kg and dMAb treated groups (data not shown).

**Table 1.** SNPs in inoculant virus comparing to A/swine/England/1353/2009

| Segment | AA_pos | NT_pos | Ref_base | Coverage | AA_change | SNP_type | A_number | T_number | G_number | C_number | N_number |
| --- | --- | --- | --- | --- | --- | --- | --- | --- | --- | --- | --- |
| KR701099.1_NA | 454 | 1361 | G | 106526 | GGT(G)->GTT(V) | nonsyn | A=38(0.000) | T=18689(0.175) | ref=G | C=38(0.000) | N=0(0.000) |
| KR701097.1_HA | 171 | 511 | A | 281680 | AAA(K)->GAA(E) | nonsyn | ref=A | T=203(0.001) | G=58387(0.207) | C=23(0.000) | N=0(0.000) |
| KR701097.1_HA | 226 | 676 | A | 104856 | AAG(K)->GAG(E) | nonsyn | ref=A | T=302(0.003) | G=39918(0.381) | C=25(0.000) | N=0(0.000) |
| KR701096.1_PA | 353 | 1059 | G | 284452 | AAG(K)->AAA(K) | syn | A=41195(0.145) | T=283(0.001) | ref=G | C=60(0.000) | N=0(0.000) |
| KR701096.1_PA | 693 | 2079 | C | 109703 | TGC(C)->TGT(C) | syn | A=218(0.002) | T=18512(0.169) | G=76(0.001) | ref=C | N=0(0.000) |

Amino acid and nucleotide numberings start at Met or ATG.

**Table 2.** Non-synonymous SNPs in day 3 samples comparing to A/swine/England/1353/2009

| Sample | Segment | AA_pos | NT_pos | Ref_base | Coverage | AA_change | SNP_type | A_number | T_number | G_number | C_number | N_number | Compare to inoculant |
| --- | --- | --- | --- | --- | --- | --- | --- | --- | --- | --- | --- | --- | --- |
| 30_naive | KR701097.1_HA | 226 | 676 | A | 271572 | AAG(K)->GAG(E) | nonsyn | ref=A | T=45(0.000) | G=236453(0.871) | C=81(0.000) | N=0(0.000) | Same as inoculant |
| 31_naive | KR701097.1_HA | 226 | 676 | A | 190623 | AAG(K)->GAG(E) | nonsyn | ref=A | T=95(0.000) | G=158242(0.830) | C=43(0.000) | N=0(0.000) | Same as inoculant |
| 51_2-12c | KR701097.1_HA | 226 | 676 | A | 8398 | AAG(K)->GAG(E) | nonsyn | ref=A | T=0(0.000) | G=8394(1.000) | C=0(0.000) | N=0(0.000) | Same as inoculant |
| 53_2-12c | KR701097.1_HA | 172 | 515 | G | 123369 | GGA(G)->GAA(E) | nonsyn | A=14937(0.121) | T=6(0.000) | ref=G | C=22(0.000) | N=0(0.000) | New SNP |
|  | KR701097.1_HA | 201 | 602 | C | 151699 | ACT(T)->ATT(I) | nonsyn | A=14(0.000) | T=16907(0.111) | G=67(0.000) | ref=C | N=0(0.000) | New SNP |
|  | KR701097.1_HA | 226 | 676 | A | 139973 | AAG(K)->GAG(E) | nonsyn | ref=A | T=37(0.000) | G=97706(0.698) | C=54(0.000) | N=0(0.000) | Same as inoculant |

Amino acid and nucleotide numberings start at Met or ATG.

## Discussion

In the last decade there has been extensive research on the use of mAbs for passive immunization against influenza. mAbs could be used as pre- or post-exposure treatment to prevent or reduce severe disease. They have the advantage of providing immediate immunity and bridging the gap between the start of a pandemic and vaccine availability. In order to test candidate mAbs and delivery platforms, we have established a reproducible and robust pig influenza challenge model and identified a protective human HA1 specific mAb, 2-12C, which can be used as a standard to benchmark other mAb candidates and delivery platforms. Both viral load and pathology were significantly reduced when the recombinant 2-12C mAb was given at 15 mg/kg intravenously 24 hours before pH1N1 challenge. Furthermore, the mAb-mediated a significant reduction in lung pathology upon administration at a lower dose as a recombinant protein (1 mg/kg) or as a dMAb (0.5 mg DNA/kg). No evidence for 2-12C driven viral evolution was detected in any group. However, it is important to reflect that 2-12C, which is a typical mAb to the globular head of HA, selects resistance mutations *in vitro* at position K130 ^31, 33^. While the epitope recognized by 2-12C has remained stable in circulating seasonal H1N1 viruses for ten years, in 2019 viruses have appeared with a substitution at N129D in the haemagglutinin that are resistant to neutralization by 2-12C (Rijal unpublished, Crick Reports September 2019, https://www.crick.ac.uk/partnerships/worldwide-influenza-centre/annual-and-interim-reports). While antibodies to the globular head are profoundly protective, they will need to be regularly updated.

Conventional mAbs remain an expensive approach from a manufacturing perspective so that a simple, cost-effective passive immunization strategy inducing sustained *in vivo* production would be extremely valuable. dMAbs, a DNA plasmid encoding mAbs have the potential to circumvent cost constraints and provide durable immunity, perhaps for the duration of an influenza pandemic or season^24–27^. Here we tested 2-12C dMAb in the pig influenza model and although the serum concentration of *in vivo* expressed dMAb was 100 times lower (∼1 μg/ml) than in pigs given the recombinant protein at 15 mg/kg (∼100 μg/ml) we observed significant protection against disease pathology in the lungs. This was encouraging as a relatively low dose of dMAb was used (0.5 mg DNA/kg), and the anti-human 2-12C response induced in the pig prevented peak concentration from being achieved (Fig. 5 and 6).

We also tested 2-12C dMAb, partially porcinised with pig Fc IgG3, and observed similar protection and an ADA response (data not shown), indicating that the human Fab still induces an antibody response. Figure 4d exemplifies a typical PK curve in the absence of an ADA. Using pig influenza-specific antibodies will circumvent this problem. We have generated a number of high affinity pig influenza neutralizing antibodies which will provide a more relevant test system to evaluate delivery of novel dMAbs and other emerging platforms in pigs. Furthermore these porcine mAbs will allow detailed investigation of pharmacokinetics and protective mechanisms in a relevant natural host large animal model.

A number of outstanding questions remain for mAb therapy and this pig model will provide an opportunity to investigate them. We still do not know the minimal protective dose of 2-12C since 15 mg/kg reduced both viral load and pathology, while 1 mg/kg reduced only pathology. In the rec 2-12C 15 mg/kg group 2-12C was detected in the BAL (320 ng/ml), in the 1 mg/kg group (8 ng/ml) and in the dMAb group (1.4 ng/ml), suggesting that it does reach mucosal surfaces although virus neutralization and reduction of BAL viral load was only observed in the 15 mg/kg group. In the 15 mg/kg group only 2-12C was detected in nasal swabs at 4 DPI, perhaps because of competition with the virus. However it remains to be determined whether reduction in viral load or gross and histo-pathology better correlate with disease morbidity and mortality. In this pig model, pH1N1 causes only a mild disease and neither moderate nor severe clinical signs were detected in any animal. However other influenza viruses, which cause more severe clinical signs in pigs would enable us to determine whether reduction of pathology without reduction in viral load correlates with symptom reduction. Studies on alternative biomarkers such as gene signatures or cytokines during mAb treatment may provide novel methods for predicting success or failure.

Here we tested the effect of prophylactic 2-12C before a high viral load is established as would be the case with therapeutic administration. However we still inoculate the pigs with a very high dose of virus and it would be important to test the effect of mAbs in a contact challenge model which more closely mimics natural infection.

In our previous studies we used the human broadly neutralizing anti-stem FI6 mAb and showed only marginal protective effect on gross pathology after aerosol delivery. We were unable to detect binding of FI6 to porcine PBMC or ADCC when porcine PBMC were used. More refined analysis of binding of human IgG to the Göttingen minipig Fc gamma receptors (FcγR) indicated that the binding affinities for human IgG to porcine FcγRIa, FcγRIIa and FcγRIIb were comparable to the respective human FcγRs^34^. However there was no binding of hu IgG1 to the poFcγRIIIa, which is an important mediator of ADCC in monocytes and natural killer cells. The lack of binding to porcine poFcγRIIIa may abrogate or greatly reduce Fc-mediated functions of human mAbs in agreement with our previous study^28^. This data suggest that there is a need for further investigation of which Fc receptors, Ig classes and subclasses are critical for therapeutic effects of mAbs *in vivo*.

An essential component in development of mAbs therapies is the need to improve existing animal models to more closely mimic humans. Unfortunately, all animal models have limitations in recapitulating the full range of disease observed in humans. Mice, guinea pigs, ferrets and non-human primates (NHP) are used for influenza-virus research, with the ferret considered to represent the “gold standard”. However mice cannot be infected with most strains of the influenza virus and do not recapitulate signs of illness observed in humans, guinea pigs do not exhibit overt signs of illness and ferrets may have different drug pharmacokinetics to humans^35–38^. NHPs share many physiological and genetic similarities to humans and are naturally susceptible to influenza virus infection, however there are ethical considerations, they are costly, not easily accessible, and generally weigh as little as 2-4 kg. In contrast, the pig is a large animal and a natural host for the same subtypes as human seasonal strains as well as a source of new human pandemic viruses^16^. Our pigs were between 11-14 kg, but pigs with equivalent weights to humans are available. A further advantage of the pig model is that the same pdmH1N1 virus circulates in both pigs and humans and has been used in human challenges as well as in our pig model ^39–41^. Therefore findings in the pig could be directly tested in humans to further validate the model.

In summary we have established a positive control protective influenza mAb and delivery method to benchmark mAb delivery platforms. This will enable testing of further improvements in the mAb delivery platforms, the effect of therapeutic administration and whether cocktails of mAbs will provide synergy by harnessing both classical neutralization and Fc mediated effector mechanisms. Furthermore, research in the pig has the additional benefit in facilitating development of new platforms for interventions in livestock diseases. We propose that the pig influenza model will become a critical tool to accelerate mAb development to the clinic by validating lead mAbs from smaller animal models and evaluating emerging delivery approaches.

## Materials and methods

### Antibody preparation

The anti-influenza human IgG1 mAb 2-12C and the anti-fluorescein human IgG1 isotype control were produced in bulk by Absolute Antibody Ltd (Redcar, UK). They were dissolved in 25mM Histidine, 150 mM NaCl, 0.02% Tween, pH6.0 diluent. We also engineered synthetic plasmid DNA to encode the human 2-12C. A single plasmid was designed to encode both the mAb 2-12C heavy and light chains under the control of a human CMV promoter and human IgG signal sequence and bovine growth hormone polyadenylation signal. The heavy and light Ig chains genes were separated by both a furin cleavage and a P2A cleavage site to ensure complete processing of the two proteins. The resulting construct was named dMAb 2-12C.

### Influenza challenge studies in pigs

All experiments were approved by the ethical review processes at the Pirbright Institute and Animal and Plant Health Agency (APHA) and conducted according to the UK Government Animal (Scientific Procedures) Act 1986. APHA conforms to ARRIVE guidelines. Two pig influenza challenge experiments were carried out.

For the first experiment, fifteen 5 weeks old Landrace x Hampshire cross, female pigs were obtained from a commercial high health status herd and were screened for absence of influenza A infection by matrix gene real time RT-PCR and for antibody-free status by hemagglutination inhibition using four swine influenza virus antigens – pdmH1N1, H1N2, H3N2 and avian-like H1N1. Pigs weighed between 11 and 14 kg (average 12.5 kg). Pigs were randomized (https://www.graphpad.com/quickcalcs/randomize1/) into three groups of five animals as follows: the first control group received 1 ml/kg histidine diluent only; the second isotype control group received 15 mg/kg anti-fluorescein IgG1 mAb intravenously and the third 2-12C group received 15mg/kg 2-12C intravenously. The antibodies were administered to the ear vein of animals sedated with stresnil. Twenty four hours after mAb administration all animals were challenged intranasally with 3 × 10^6^ PFU of pandemic swine H1N1 isolate, A/swine/England/1353/2009 (pH1N1) in 4 ml (2ml per nostril) using a mucosal atomization device MAD300 (Wolfe Tory Medical).

In the second challenge experiment thirty, 5 weeks old Landrace x Hampshire cross, influenza free female pigs were obtained and their influenza free status confirmed as above. The weights of the pigs were between 12 and 14 kg (average of 12.9 kg). The pigs were randomized into 4 groups of 6 as follows: control untreated; 15 mg/kg recombinant 2-12C intravenously; 1 mg/kg recombinant 2-12C intravenously and the final group received 6 mg human dMAb 2-12C. Recombinant 2-12C protein was administered intravenously 24 hours before the pH1N1 challenge to sedated pigs and dMAbs were administered by electroporation to pigs sedated with 4mg/kg Zoletil and 0.04 mg/kg Domitor, 6 days before the pH1N1 influenza challenge. Six mg of dMAb 2-12C pDNA was formulated with 135 U/ml Hylenex (Halozyme, CA), the formulation was administered intramuscularly to the back left quadriceps of the animal, and followed by *in vivo* electroporation using the CELLECTRA® constant current device. All animals were challenged with 3.4 × 10^6^ PFU pH1N1 in 4ml (2ml per nostril) using MAD. Clinical signs (temperature, state of breathing, coughing, nasal discharge, appetite, altered behavior) observed were mild and none of the pigs developed moderate or severe disease.

### Pathological and histopathological examination of lungs

Animals were humanely killed four days post infection (DPI) with an overdose of pentobarbital sodium anesthetic. The lungs were removed and digital photographs taken of the dorsal and ventral aspects. Macroscopic pathology was scored blind as previously reported^42^. The percentage of the lung displaying gross lesions for each animal was calculated using image analysis software (Fiji ImageJ) on the digital photographs. Lung tissue samples from cranial, middle and caudal lung lobes were taken from the left lung and collected into 10% neutral buffered formalin for routine histological processing. Formalin fixed tissues were paraffin wax-embedded and 4-µm sections cut and routinely stained with hematoxylin and eosin (H&E). Immunohistochemical detection of influenza A virus nucleoprotein was performed in 4-µm tissue sections as previously described^43^. Histopathological changes in the stained lung tissue sections were scored by a veterinary pathologist blinded to the treatment group. Lung histopathology was scored using five parameters (necrosis of the bronchiolar epithelium, airway inflammation, perivascular/bronchiolar cuffing, alveolar exudates and septal inflammation) scored on a 5-point scale of 0 to 4 and then summed to give a total slide score ranging from 0 to 20 per slide and a total animal score from 0 to 60^44^. The slides were also scored using the “Iowa” method, that also takes into account the amount of viral antigen present in the sample, as described^45^.

### Tissue sample processing

Four nasal swabs (NS) (two per nostril) were taken at 0, 1, 2, 3, 4 DPI. The swabs were placed into 1 ml of Trizol or 2 ml of virus transport medium comprising tissue culture medium 199 (Sigma-Aldrich) supplemented with 25mM HEPES, 0.035% sodium bicarbonate, 0.5% bovine serum albumin (BSA), penicillin 100 IU/ml, streptomycin 100 µg/ml and nystatin 0.25 µg/ml, vortexed, centrifuged to remove debris and stored at −80°C for subsequent virus titration. Blood samples were collected at the start of the study (prior to antibody administration) and at the indicated times post mAb delivery and challenge. Broncho-alveolar lavage (BAL) was collected from the entire left lung with 150ml of virus transport medium (described above). BAL samples were centrifuged at 300 x g for 15 minutes, supernatant was removed, aliquoted and frozen for antibody analysis.

### Virus titration and viral RNA isolation

Viral titers in nasal swabs, BAL and accessory lung lobe were determined by plaque assay on MDCK cells (Central Service Unit, The Pirbright Institute, UK). Samples were 10-fold serially diluted in Dulbecco’s Modified Eagle’s Medium (DMEM) and 100µl overlayed on confluent MDCK cells in 12 well tissue culture plates. After 1 hour, the plates were washed and overlayed with 2ml of 0.66 % agarose containing culture medium. Plates were incubated at 37°C for 48 to 72 hours and plaques visualized using 0.1% crystal violet. For sequencing nasal swabs were collected in 1 ml of Trizol (Invitrogen, Thermo Fisher Scientific, UK) and RNA was extracted by chloroform and isopropanol precipitation.

### Next generation sequencing

Viral RNA enriched library production and viral cDNA library sequencing were as previously reported^46^. Briefly, isolated RNA was amplified using the Ovation RNA-Seq system V2 from NuGEN (NuGEN, San Carlos, CA). The amplified total cDNAs were analyzed by an Agilent 2100 Bioanalyzer using the Agilent High Sensitivity DNA Kit (Agilent Technologies, Santa Clara, CA) and sheared to 150bp on the Covaris S2 machine (Covaris, Woburn, MA). Approximately 400 ng of amplified cDNA was used to generate the Illumina sequencing library using the Agilent SureSelectXT Target Enrichment Kit (Agilent, Santa Clara, CA) for Illumina Multiplex Sequencing by using enrichment probes designed for A/California/04/2009(H1N1) virus. Enriched Illumina sequencing libraries were sequenced on an Illumina NextSeq sequencer (Illumina, San Diego, CA).

### Data analysis

Reads were mapped to the HISAT2 indexed A/swine/England/1353/2009 (pH1N1) genome using HISAT2 (release 2.0.5, https://ccb.jhu.edu/software/hisat2/index.shtml) downloaded from the Center for Computational Biology, Johns Hopkins University^47^. SAMtools mpileup (version 2.1.0)^48^ was used to make SNP calls with minimum base Phred quality score as 25. A reported SNP call was one that satisfied the following criteria at the SNP position: 1) more than 100 reads at that position^49–51^ 2) reads present from both directions, 3) variant calls exactly at the end of the read eliminated and 4) reads with bases that are different to reference more than 10% of the aligned reads.

### Microneutralization assay

Neutralizing Ab titers were determined in serum and BAL fluid using a microneutralization (MN) assay as previously described^52^. In brief, pig sera or BAL fluid were heat-treated for 30 min at 56°C and diluted 1:10 for serum and 1:2 for BAL as a starting point for the assay. Fifty µl of serially diluted samples were incubated with an equal volume of pH1N1 (the virus was titered beforehand in the absence of serum to determine the PFU/ml necessary to yield a plateau infection in the MN assay). After 2h MDCK SIAT-1 cells at 3 × 10^4^ cells/well were added to the serum/virus and incubated for 18 h. The fixed and permeabilized cell monolayer was stained with anti-nucleoprotein (Clone: AA5H, Bio-Rad Antibodies, UK) followed by goat anti-mouse HRP (DAKO) antibody. After addition of the 3,3’,5,5’-tetramethylbenzidine (TMB) substrate the reaction was stopped with 1M sulfuric acid and absorbance was measured at 450 nm and 570 nm (reference wavelength) on the Cytation3 Imaging Reader (Biotek). The MN titers were expressed as half maximal inhibitory dilution (50% Inhibitory titer is: midpoint between uninfected control wells and virus-infected positive controls) derived by linear interpolation from neighbouring points in the titration curve.

### Enzyme-linked immunosorbent assays (ELISA)

Antibody titers against the pH1N1 HA in the serum, BAL fluid and nasal swabs were determined by ELISA. The recombinant HA protein of A/Eng/195/2009 containing a C-terminal thrombin cleavage site, a trimerization sequence, a hexahistidine tag and a BirA recognition sequence was expressed in HEK293 cells and purified as described previously^52^. Ninety six-well microtiter plates (Maxi Sorp, Nunc, Sigma-Aldrich, UK) were coated with 50 µL recombinant protein at a concentration of 1 µg/mL in carbonate buffer overnight at 4 °C. Plates were blocked with 200 μl blocking solution of 4% milk powder in PBS, supplemented with 0.05% Tween-20 (PBS-T) for 2 h at room temperature. Samples were serially diluted in PBS-T with 4% milk powder and added to the wells for 1h on a rocking platform. The plates were washed three times with PBS-T and 100 μl of horseradish peroxidase (HRP)-conjugated goat anti-human or goat anti-pig Fc fragment secondary antibody (Bethyl Laboratories) diluted in PBS-T with 4% milk powder was added and plates incubated for 1 h at room temperature. The plates were washed four times with PBS-T and developed with 100 µL/well TMB High Sensitivity substrate solution (BioLegend, UK). After 5 to 10 min the reaction was stopped with 100 µL 1M sulfuric acid and the plates were read at 450 and 570 nm with the Cytation3 Imaging Reader (Biotek). A standard curve was generated using rec 2-12C antibody (Absolute Antibody). The data were analyzed in Microsoft Excel and GraphPad Prism. The cut off value was defined as the average of all blank wells plus three times the standard deviation of the blank wells.

For detection of an antibody response to 2-12C in pigs (ADA), ninety six-well microtiter plates (Maxi Sorp, Nunc, Sigma-Aldrich) were coated with 2 μg/ml rec 2-12C (Absolute Antibody) in PBS-T overnight at 4°C. The plates were washed and blocked. Pre-diluted serum samples in T-PBS were added and incubated for 2h at room temperature. The plates were washed 3 times and goat anti-pig H+L-HRP (Bethyl Laboratories) was added for 1 h at room temperature and developed as above.

### *In vitro* transfection of dMAb 2-12C

Lipofectamine 3000 transfection kit (ThermoFisher, Waltham, MA) was used to transfect adherent HEK 293T cells (ATCC® CRL11268™) with dAMb 2-12C plasmids. Medium was harvested 72 h post transfection and filtered using 0.22 µm stericup-GP vacuum filtration system (Millipore, Burlington, MA). The supernatant from pDNA transfected cells was purified using Protein G GraviTrap (GE Healthcare, Chicago, IL) according to the manufacturer’s instructions. The eluted protein was concentrated by Amicon Ultra-15 Centrifugal Filter unit (30 kDa) and quantified by nanodrop.

### Western blot

0.5 µg of each sample was loaded on a NuPAGE™ 4-12% Bis-Tris gel (ThermoFisher). Precision Plus Protein Kaleidoscope Prestained Protein Standard (Bio-Rad, Hercules, CA) was used as the standard marker. The gel was transferred to a PVDF membrane using iBlot™ 2 Transfer device (Invitrogen, Waltham, MA). The membrane was blocked with goat histology buffer (1% BSA (Sigma, St. Louis, MO), 2% goat serum (Sigma), 0.3% Triton-X (Sigma) and 0.025% 1g/ml Sodium Azide (Sigma) in PBS) for 30 minutes at room temperature. Goat anti-human IgG-Fc fragment antibody (A80-104A, Bethyl, Montgomery, TX) diluted in 1:1000 in goat histology buffer was added and incubated for 1 hour at room temperature. After washing the blot for 5 minutes in DPBS (HyClone, Logan, UT) three times, donkey anti-goat IgG HRP antibody (Abcam, Cambridge, UK) in 1:2000 dilution in goat histology buffer was added and incubated for 1 hour at room temperature. After washing the blot three times for 5 minutes in DPBS, the membrane was developed using ECL^™^ Prime Western Blotting system (GE Healthcare) and imaged using Protein Simple FluorChem System.

### *In vivo* dMAb 2-12C mouse and pig immunogenicity and pharmacokinetic studies

Female BALB/c mice between 4 and 6 weeks of age were group-housed with *ad libitum* access to feed and water. Husbandry was provided by Acculab (San Diego, CA) and all procedures were in compliance with the standards and protocols of the Institutional Animal Care and Use Committee (IACUC) at Acculab. To deplete T cell populations BALB/c mice received one intraperitoneal injection of 500 µg of anti-mouse CD4 (BE0003-1, BioXCell, West Lebanon, NH) and 500 µg of anti-mouse CD8α (BioXCell, BE0117) in 300 µl of PBS. 200 µg dMAb plasmid DNA was formulated with 135 U/ml of human recombinant hyaluronidase (Hylenex®, Halozyme, San Diego, CA). Mice received an intramuscular (IM) administration of formulation followed by electroporation (EP). EP was delivered at the injection site with the CELLECTRA-3P® adaptive *in vivo* electroporation system. An array of three needle electrodes with 3 mm insertion depth was used. The EP treatment consists of two sets of pulses with 0.1 Amp constant current with the second pulse set delayed by 3 seconds. Within each set there are two 52 ms pulses with a 198 ms delay between the pulses. 100 µl recombinant 2-12C was injected intravenously (IV) at a dose of 1 mg/kg.

Pigs received intramuscular administration of up to 24 mg of dMAb plasmid DNA formulated with 135 U/ml of human recombinant hyaluronidase. EP was delivered at the injection site with the CELLECTRA-5P® adaptive *in vivo* electroporation system. For pharmacokinetic (PK) studies, blood samples were drawn at the indicated time points.

### Immunofluorescence staining and imaging

*In vivo* expressed 2-12C: 3 days after IM delivery of dMAb 2-12C pDNA, mouse and pig muscle tissues were harvested, fixed in 10% Neutral-buffered Formalin (BBC Biochemical, Stanford, MA) and immersed in 30% sucrose (Sigma) in deionised water for *in vivo* staining of dMAb expression. Tissues were then embedded into O.C.T. compound (Sakura Finetek, Torrance, CA) and snap-frozen. Frozen tissue blocks were sectioned to a thickness of 18 µm. Slides were incubated with Blocking-Buffer (0.3% Triton-X (Sigma), 2% donkey serum in PBS) for 30 min, covered with Parafilm. Goat anti-human IgG-Fc antibody (Bethyl) was diluted 1:100 in incubation buffer (1% BSA (Sigma), 2% donkey serum, 0.3% Triton-X (Sigma) and 0.025% 1g/ml Sodium Azide (Sigma) in PBS). 150 µl of staining solution was added to each slide and incubated for 2 hrs. Slides were washed in PBS three times. Donkey anti-goat IgG AF488 (Abcam, Cambridge, UK) was diluted 1:200 in incubation buffer and 50 µl was added to each section. Slides were washed after 1hr incubation and mounted with DAPI-Fluoromount (SouthernBiotech, Birmingham, AL) and covered. Slides were imaged with a BX51 Fluorescent microscope (Olympus, Center Valley, PA) equipped with Retiga3000 monochromatic camera (QImaging, Surrey, Canada).

### Quantification of human IgG in dMAb 2-12C treated animals

96-well assay plates (Thermo Scientific) were coated with 1 µg/well goat anti-human IgG-Fc fragment antibody (Bethyl) in DPBS (ThermoFisher) overnight at 4°C. Next day plates were washed with 0.2%Tween-20 in PBS and blocked with 10% FBS in DPBS for 1hr at room temperature. The serum samples were diluted in 1% FBS in 0.2% Tween-PBS and 100 µl of this mix was added to the washed assay plate. Additionally, a standard curve of recombinant 2-12C mAb prepared as 1:2 serial dilutions, starting at 500 ng/ml in dilution buffer and added in duplicate to each assay plate. Samples and standard were incubated for 1hr at room temperature. After washing, the plates were incubated with a 1:10,000 dilution of goat anti-human IgG-Fc Antibody HRP (Bethyl, A80-104P) for 1hr at room temperature. For detection SureBlue Substrate solution (Seracare, Milford, MA) was added to the washed plates. The reaction was stopped by adding TMB Stop Solution (Seracare, Milford, MA) after 6min to the assay plates. The optical density (O.D.) were read at 450nm. The serum concentration was interpolated from the standard curve using a sigmoidal four parameter logistic curve fit for log of the concentration.

### Statistical analysis

One-way non-parametric ANOVA (Kruskall-Wallis) with Dunn’s post test for multiple comparisons was performed using GraphPad Prism 8.3.

## Acknowledgements

We are grateful to the animal staff at APHA for excellent animal care and for providing the challenge swine A/Sw/Eng/ 1353/09 influenza virus strain (DEFRA SwIV surveillance programme SW3401).We thank Carlo Bianco from APHA for his support in the image analysis. This work was supported by Bill & Melinda Gates Foundation Grant OPP1201470 and Biotechnology and Biological Sciences Research Council (BBSRC) Grant BBS/E/I/ 00007031. This work was supported in part by the Intramural Research Program of the National Institute of Allergy and Infectious Diseases, National Institutes of Health (YX, JCK, JKT).

## Author contributions

ET, AM, BH, KB, DW, TS conceived, designed and coordinated the study. ET, AM, BH, BC, EB, GG, HB, KS, SR, PF, AP and SE performed animal experiments, processed samples and analyzed the data. VM, TC, AN and MB carried out post mortem and pathological analyses. YX, JK and JT performed sequencing analysis. ET, AM, PB, TS, KB wrote and revised the manuscript and figures. AT and PR provided reagents and performed microneutralization assays.

